# Wildlife damage to tropical agroforests: predicting primate-induced fruit loss using species’ energy requirements

**DOI:** 10.64898/2026.10.21.760699

**Authors:** Bruno Simmen, Guillaume Correa-Pimpao, Romane Ibanez, Jeanne Tonnabel, Laurent Tarnaud

## Abstract

1. Quantitatively assessing wildlife damage to food crops in tropical agroforestry systems is notoriously problematic. At the same time, the animal species likely to destroy these crops, such as primates, are often endangered and a source of conflict. Resolving these conflicts may require the implementation of financial compensation where non-monetary solutions are unavailable. Currently, however, there is no suitable method for assessing fruit losses due to the complexities of agroforestry plots used by most small households practising polyculture-based agriculture.

2. We constructed an energy balance model to predict the quantity of cultivated fruit and other locally sourced food required by crop-dependent primates to meet their metabolic needs. The proposed method is based on the principle that, under conditions of energy balance, an animal’s daily energy intake corresponds to its total energy expenditure. The ecophysiological model incorporated the daily food and energy intake of wild primates, measured using the bite-size method and the nutritional composition of food, as well as their total energy expenditure and body composition, measured using doubly labelled water. We tested the model’s predictions on lemurs foraging in several agroforests in Mayotte, and on two frugivorous primate species in Madagascar during different seasons. Additionally, we evaluated whether more conventional dietary analysis methods (feeding-time sampling), or published energetics data could reliably inform the model, and reduce the costs and constraints of field observations.

3. The predictions were generally consistent with direct estimates of food intake (91% ± 25%, n = 14 across different seasons), though this was influenced by the lemurs’ evolutionary-driven seasonal fattening strategy. Time-derived estimates using correction factors or published measurements of total energy expenditure were also consistent, but further studies are required to confirm these findings.

4. *Policy implications*. The study provides proof of concept for a promising approach to quantifying wildlife-related agricultural losses in small-scale agroforestry. This innovative method provides policymakers, farmers’ associations, and non-governmental organisations with sufficiently accurate quantitative estimates to support the development of evidence-based financial compensation schemes.

## 1 Introduction

Conflicts between humans and wildlife involving damage to food crops have been widely documented (FAO, 2026; IUCN, 2023). In low-income tropical countries, most small-scale farming families own small plots of land to grow a variety of edible plants, while a few others cultivate plant resources as a cash crop, which can be their main source of income. Awareness of the frequent deleterious effect of human-wildlife conflicts on biodiversity as well as the destruction of habitats for new agricultural activities has led to the development of various action plans aimed at reducing poverty while maintaining or restoring ecosystem function.

Since decades, compensation schemes to help farmers and promote conservation, such as conditional payments for environmental services (PES), have been used in many areas of the world (Jayachandran et al., 2017; Pagiola et al., 2005; Wunder et al., 2020). However, implementing PES in developing countries is fraught with difficulties, for example when it relies on weak institutions (Pattanayak et al., 2010). In fact, even high-income countries with tropical regions containing agroforests and other agrobiodiversity systems face practical and political difficulties in implementing action plans. Financial compensation for small-scale crop production may be refused by local authorities simply because of the lack of methods for satisfactorily estimating losses in fruit and vegetable production (e.g. Gonçalves et al., 2020; Hill, 2018). Indeed, unlike stem damage, the damage caused to this type of crop production by wildlife is far from being reliably quantified (Ango et al., 2017; Conforti et al., 2020; Naughton-Treves, 1997; Wallace & Hill, 2012). This applies to both small farming households practising polyculture-based agriculture at a subsistence level and those to some extent integrating the market economy. Currently, there are no methods adapted to complex tropical agroforestry systems, where cultivation practices involve mixing various edible herbaceous plants, trees and lianas in spatially superimposed crops with staggered fruiting periods on small plots (Schroth et al., 2004). What’s more, the conservation status of endangered animal species that can destroy crops but play a major role in forest regeneration through zoochorous dispersal systems, for example primates or elephants among large mammals, requires the implementation of gentle solutions that are accepted by the different socio-economic actors.

The aim of this study was to design a method that is sufficiently accurate for assessing fruit crop losses in mixed-crop agroforestry systems. We chose primates as our case study because most of these species survive outside protected areas and are threatened or critically endangered, creating acute problems in resolving conflicts between small farmers and wildlife (Estrada et al., 2012, 2017; FAO, 2026; Schwitzer et al., 2014; Torres-Romero et al., 2023). It is therefore crucial to develop quantitative methods that demonstrate the feasibility of the approach to stakeholders. Since it is rarely possible to directly quantify the amount of fruit consumed by wild animals in such agroforests, we propose an ecophysiology-driven approach to estimating dietary intake. The general principle is to predict how much of each known food a primate needs to consume daily to meet its total daily energy requirements. This can then be used to estimate the quantity of fruit and other vegetation consumed by groups foraging in the area. This method requires the composition of the diet to be identified beforehand, at least the staple foods, which is relatively straightforward when animals consistently use crops. We primarily built our model on brown lemurs (*Eulemur fulvus*, Strepsirrhini: Lemuridae) living on Mayotte Island. We determined their diet, total energy expenditure (TEE) and nutrient intake across different seasons in order to test the effectiveness of the model. From an operational point of view, we compared the consumption values measured using the mouthful method for the different foods ingested by the individuals we observed continuously from dawn to dusk, with the consumption values predicted from their TEE. This prediction is basically informed by the relative proportions of foods in the diet, as well as the metabolizable energy content of the foodstuffs. Since this approach is based on the working hypothesis that the animals achieve a balance between energy intake and expenditure, we also measured their body mass and composition.

Given the challenges associated with quantitative feeding ecology studies, and the fact that primates impact many cultivated areas around the tropical world, we explored alternative techniques to make the method widely applicable. In particular, continuous recording of feeding activity is not always possible in the field, so we investigated whether conventional time-sampling methods and varying levels of knowledge about diets could reliably estimate daily food intake. Finally, since measuring energy expenditure can be challenging and costly, we examined whether published mass-TEE data could accurately predict daily food intake for a few frugivorous species across different locations and years. We tested this in brown lemurs in a different year or study site in Mayotte, as well as in other primates in Madagascar (*Eulemur rufifrons x collaris* and *Lemur catta*, Strepsirrhine, Lemuridae). Overall, the findings of this study demonstrate that prediction-based estimates are accurate enough to inform the implementation of financial solutions. We discuss the extent to which the observed variability is due to the approximations inherent in nutritional ecology approaches and energy strategies involving seasonal fattening.

## 2 Materials and methods

### 2.1 Study site and season

Following a preliminary survey conducted in Mayotte in October 2021, we selected two agroforestry plots for further investigation. Our selection was based on the typology of the agricultural system (i.e. polycultures dedicated to food or cash crops and orchards), the presence of numerous lemurs, the possibility of continuously observing the animals, the feasibility of capturing and recapturing lemurs in order to carry out the energy physiology experiment, and the landowners’ agreement. The study took place at both sites in parallel during two seasons in 2022, between February 15 and May 15, and between June 20 and July 29. One study site was located in Tsararano (12°49’53’’S, 45°10’22.8’’E), an irrigated agroforestry plot spanning several hectares dedicated to local fruit and transformed product trade (e.g., fruit juices). The second site, located in Kahany (12°49’25.5’’S, 45°08’12.1’’E), is a small-scale agroforest dedicated to family consumption of fruit and vegetables, a model that is prevalent throughout Mayotte island (Agreste, 2022a). Thirteen relatively homogeneous zones are recognised in Tsararano, where perennial cultivated trees (mango, guava, Cythera prune, breadfruit, avocado, edible Annonaceae, *Albizia*, etc.) and native trees and vines (e.g. *Saba comorensis*) coexist with small banana plantations (Laurane Rami, personal communication). In open areas, passion fruit, pineapple, papaya, and a few other edible plants are grown (27 species over 4.8ha, 119 trees >10cm diameter per ha, basal area: 11.7 m^2^ ha^-1^).

In Kahany, the wooded area is essentially a system of intertwined polycultures with no dominant culture, but which can nevertheless be physiognomically categorized into citrus, cocoa-coconut-banana and litchi-bilimbi zones. There, trees and edible lianas produce mangoes, bananas, breadfruit, citrus, coconuts, cocoa, litchees, pepper and Indian laurel fruits among other plant species like papaya, coffee and vanilla (28 species over 1 ha, 150 trees >10cm diameter per ha, basal area: 14.2 m^2^ ha^-1^; L. Rami, personal communication). Both sites are connected to a close protected departmental forest reserve by secondary forest corridors and other agricultural plots with substantial forest cover. Fruit was available throughout the study period, but the types of fruit available changed between the wet and dry seasons (Fig. S1).

### 2.2 Lemur focal groups

In the two study sites, lemur groups are almost entirely dependent on the food plants available in the plots. Their home ranges overlap slightly and vary between 1 and 4 hectares (according to collar-mounted GPS and manual GPS recording performed during the observation study; Simmen et al., 2024; this study). We counted a total of 63 lemurs (five groups) in Tsararano and 42 lemurs (four groups) in Kahany, including 10 infants. Sub-adults cannot be distinguished by size from adults (i.e., over 3 years old; unpublished data, Laurent Tarnaud). We conducted the behavioural study during part of the rainy and dry seasons, focusing on four contiguous groups in Tsararano and two contiguous groups in Kahany. Reproduction is highly seasonal in this species, with a birth peak in October and weaning around March (Tarnaud, 2004). Focal females were not lactating during the study, but some females may have been in very early gestation during the dry months of the study.

### 2.3 Diet composition, food intake and feeding duration

#### 2.3.1 Quantitative determination of diet

The excellent visibility and permanence of groups well accustomed to human observers enabled us to follow focal groups from dawn to dusk for over 300 h during the study (29 full days balanced across groups in each season and at each site). To determine the composition of the diet, one observer used the animal focal sampling method (Altmann, 1974) to count the number of whole foods (e.g. berries) or bites consumed by a particular individual within a group, or the number of mouthfuls devoted to each food, over a period of 30 min (Hladik 1977, 2011; Rothman et al., 2012). High-resolution binoculars were used when necessary. The different parts of fruits, leaves, and flowers eaten were weighed thereafter by sampling food (> 10 samples per food item) from the feeding trees whenever possible, and using a portable balance (200 ± 0.1g). Observations alternated between adult/subadult members of the group during an observation day and between groups in different days. All digestible (e.g., fruit pulp) and indigestible parts (e.g., mature seeds, which passed undigested through the intestinal tract) of the food swallowed were weighed. During mouthful counts, the corresponding feeding duration (in minutes) was recorded, enabling us to calculate feeding rates (g min^-1^) and use them when the focal animal was less visible during a feeding episode or for other calculations as below.

Like some other cathemeral species, *Eulemur fulvus* exhibits nocturnal activity in addition to its daytime activity, mostly around the time of the full moon (Donati et al., 2009; Kappeler & Erkert, 2003). In Mayotte, accelerometers fitted to members of four focal lemur groups in parallel with the feeding ecology survey (Tsararano) revealed that lemurs were far more active during the day than at night throughout the study (Simmen et al., 2024; Simmen, personal communication). Additionally, quantitative assessments of nocturnal food consumption in *Eulemur* (Simmen et al., 2007; Tarnaud, 2006) suggest that this would represent only a small proportion of the lemurs’ daily intake (e.g. <10% in *E. fulvus* in Mayotte, all moon phases combined; Tarnaud, 2006).

#### 2.3.2 Alternative method for assessing relative dietary plant proportions

We employed a second method to estimate food proportions in the diet. The aim was to establish whether feeding data obtained using a more conventional, albeit less direct, method could be reliably incorporated into the TEE-based predictive models developed further (see below). To this end, a second observer conducted instantaneous and scan sampling (Altmann, 1974) of all visible group members simultaneously with observer 1. The scans were repeated at 5-minute intervals (total: 3410 scans). We assigned a score of one unit to each food item eaten by each visible animal during a scan period. Because time-sampling methods do not properly reflect the proportions of food actually ingested (Hladik, 1977; Visalberghi & Addessi, 2003; Zinner, 1999), we weighted the feeding time data by the ingestion rate of each food (i.e., the amount of food ingested per unit of time; Nakagawa, 2009; Rothman et al., 2012).

### 2.4 Total energy expenditure (TEE) and body composition

The metabolic measurements and methodological details for determining total daily energy expenditure and body composition in this lemur population are derived from Simmen et al.’s (2024) paper. These were measured simultaneously with the present study on the same focal groups during two 10-day sessions in early March and late June/early July 2022. As the body mass, body composition, and mass-specific energy expenditure of Kahany lemurs were in the same range as those of lemurs captured in Tsararano, we pooled TEE and body mass data to obtain average values for each season and to use these reference values in further calculation. Daily energy expenditure was 557 ± 144 kJ d^-1^ (n = 10) for individuals weighing 2.01 ± 0.28 kg during the wet season and 481 ± 68 kJ d^-1^ (n = 3) for individuals weighing 1.70 ± 0.10 kg during the dry season. According to the isotopic analysis and skinfold thickness measurements, animals caught during the dry season had a lower fat mass than those caught during the wet season (Simmen et al., 2024; Simmen, personal communication).

### 2.5 Data processing

#### 2.5.1 Daily food intake and fruit crop losses

Based on the focal animal observations (mouthful method), we calculated the average daily food intake (in g day^-1^ per individual) for each study site and season. For a given observation day, we summed the amount of food consumed by focal individuals during their successive 30-minute feeding sequences, and averaged this daily value over the different observation days. By weighing the food parts consumed and not consumed (e.g., pulp, fibrous mesocarp, peduncles, seeds…), we reconstructed the average weight of intact fruits (and other foods) handled per lemur per day in Tsararano and Kahany. The results were extrapolated to the population level, based on the complete lemur census at each site. Lemur developmental stages were classified as adult/subadult, juvenile and infant. Sub-adults cannot be distinguished from adults by size (i.e. those over three years old; unpublished data, Laurent Tarnaud), and there is no difference in body weight between the sexes (Simmen et al., 2024; Tarnaud, 2024). We assumed that the body mass and energy requirements of immatures corresponded to 30% of those of adult lemurs.

#### 2.5.2 Predictive estimates of food intake (Predictions 1 and 2) and food crop losses

To predict the amount of food needed to meet energy requirements, we calculated the contribution of each food to the total daily energy needs as follows: for each food item, we multiplied its contribution to total food intake (%), as derived from focal observation (see §2.5.1), by its digestible energy content (see below). The results, in kJ g^-1^, were adjusted to the average TEE (kJ d^-1^) measured (see §2.4), maintaining proportionality between the different food items to ensure the total energy input equalled the TEE. We then back-calculated the quantity (in g) of each food that should be consumed, and consequently the predicted total daily food intake (Prediction 1). An alternative predictive calculation of the daily food intake required to meet TEE (Prediction 2) was performed using the relative food proportions determined by time-sampling scores, as described above.

We derived the digestible energy content of tropical fruits and other plant parts from the reference tables of the FAO, supplemented with data from other databases, and specific publications (Table S1). We performed additional macronutrient analyses on the only native plant sample with a high ranking in the diet (*Saba comorensis*), corrected by published digestibility measurements (fibre digestibility: 0.41% for low-fibre food in *E. fulvus*; Campbell et al., 2004a). The food’s energy content was calculated (in kJ g^-1^) by multiplying the kilocalories by 4.18 (Rothman et al., 2012). In Kahany, the proportion of the diet for which energy data was available was > 92% in the wet and dry seasons. In Tsararano, the proportion was >96%. For the few remaining food items that had not been analysed, we used the median energy value from comparable categories of vegetative or reproductive plant parts to estimate total energy intake as closely as possible.

Fruit crop losses were then calculated from Predictions 1 and 2 as described in §2.5.1.

#### 2.5.3 Testing the energetics-based method in other primates

We applied the energetics-based method to four other primate populations using TEE measurements that were here collected independently from the diet study. These case studies were chosen because, to our knowledge, they are the only frugivorous primates for which quantitative data on both food intake and energy expenditure are available. The first case study focused on two populations of *Eulemur fulvus*: one that was surveyed in the same location of Kahany as previously, but in a different year (hereafter referred to as ‘Kahany2’); and one that was surveyed in an anthropogenic area including orchards in Passamainty (12°48’5.86“S, 45°12’16.57”E), Mayotte. Daily food intake was recorded at both sites during the wet and dry seasons of 2008. A second case study examined two species of frugivorous-folivorous lemurs that coexist in a gallery forest adjacent to open, moderately populated areas in southern Madagascar (Malaza in the Berenty Private Reserve; 25°0.29’S, 46°19.37’E): a hybrid population of brown lemurs (*Eulemur rufifrons × collaris*) and a population of ring-tailed lemurs (*Lemur catta*). These lemurs are medium-sized primates with relatively similar body mass and a highly seasonal reproductive cycle with similar timing and duration patterns. Food consumption was estimated during three periods of the year: the dry and early wet seasons of 1998, and the late wet season of 1999 (Simmen et al., 2003). The diets and energy intake of the two species, including both natural and anthropogenic foods, were determined simultaneously across the three seasons. Their TEE and body composition were measured during the late wet season of 2006 (Simmen et al., 2010).

The daily food intake and relative proportions of food items in the diet were assessed in each case study using the mouthful approach and focal animal sampling over continuous observation periods (Simmen et al., 2003). On average, the set of food items used to predict daily food intake based on energy requirements of all these lemurs accounted for 93% of the total diet (range: 83–100% across 10 study cases, including seasonal variations). The nutrient and energy content of the diets was also determined (Simmen & Rasamimanana, 2018; this study).

As the dietary and TEE studies covered different time periods (several seasons for diet and one season for TEE), the dietary intake predictions for the different seasons were based on a single TEE data point. In line with the ‘constrained energy model’ principle (Pontzer, 2015; Pontzer et al., 2014; Pontzer & Trexler 2026), which posits that TEE only fluctuates within a narrow physiological range, we reasoned that a single TEE data point could be used as a reference when testing our model. The concept of constrained energy expenditure has been supported by meta-analyses of energy expenditure in humans, non-human primates, and other placental mammals, as well as by studies conducted across different seasons or between captive and wild individuals of the same species (Nagy et al., 1999; Pontzer, 2015; Pontzer et al., 2014; Simmen et al., 2024). Under habitual physical activity conditions, energy expenditure is primarily dependent on body mass, particularly fat-free mass (Pontzer et al., 2016, 2021).

## 3 Results

### 3.1 Tsararano and Kahany lemurs

#### 3.1.1 Diet composition

In Tsararano, the diet consisted of 27–28 foods during the wet and dry seasons, respectively, of which more than 78% were of anthropogenic origin. In Kahany, 22 and 18 items made up the diet, of which more than 86% were anthropogenic foods. Diet diversity was low since only three to five food items accounted for more than 80% of the ingested matter in each site and season. Fruit was the main food category whatever the study site or season. Although the set of most important foods derived from continuous consumption data did not differ from that determined from the duration of feeding sequences recorded simultaneously on the same focal animal (30-min focal sampling), the relative proportions of these foods varied considerably (see Table S2). This was due to the high ingestion rates of certain items, as in the case of banana pulp or papaya, which were disproportionately represented in terms of mass consumed (e.g., Tsararano dry, Kahany dry).

As with the focal animal sampling results described above, the set of main foods determined by the scan sampling method sometimes differed considerably, depending on whether the food units were weighted by intake rates or not (e.g. papaya in Kahany dry season; Table S3).

#### 3.1.2 Daily food intake and estimates based on continuous recording of mouthfuls (Prediction 1)

Tsararano and Kahany lemurs consumed more food during the warm, rainy austral summer than during the dry austral winter (season: *F*(1, 26) = 5.90, *p* < 0.05, site: *F*(1, 26) = 1.33, *p* < 0.1), which was consistent with seasonal changes in body composition (Methods section, §2.4). There was no significant effect of the season x site interaction term (*F*(1,25) = 3.292, *p* < 0.1) (two-way analysis of variance, using the R packages rstatix (Kassambara, 2026) and performance (Lüdecke et al., 2021), in R version 4.5.2 (R Core Team, 2025) and RStudio version 2024.12.1+563).

After converting ingested fresh material into digestible material (e.g. fruit pulp, minus the swallowed seeds) and metabolizable energy intake, we found that lemurs ingested far more energy than they needed to meet their total energy requirements during the rainy season, in contrast to the pattern observed during the dry season. Compared with the TEE-based prediction (Prediction 1), the estimated daily consumption of edible parts (i.e. excluding fruit seeds) based on direct observation was 52% higher at Tsararano (354 vs 233 g d⁻¹) and 38% higher at Kahany (249 vs 181 g d⁻¹) during the wet season (Table 1; Table S4 for the formulas and results). During the dry season, a more balanced energy input/output was observed than in summer. The estimated consumption during the dry months was 3% (176 g d⁻¹) higher than the predicted consumption at Tsararano, and 14% higher (225 g d⁻¹) at Kahany.

**Table 1:**
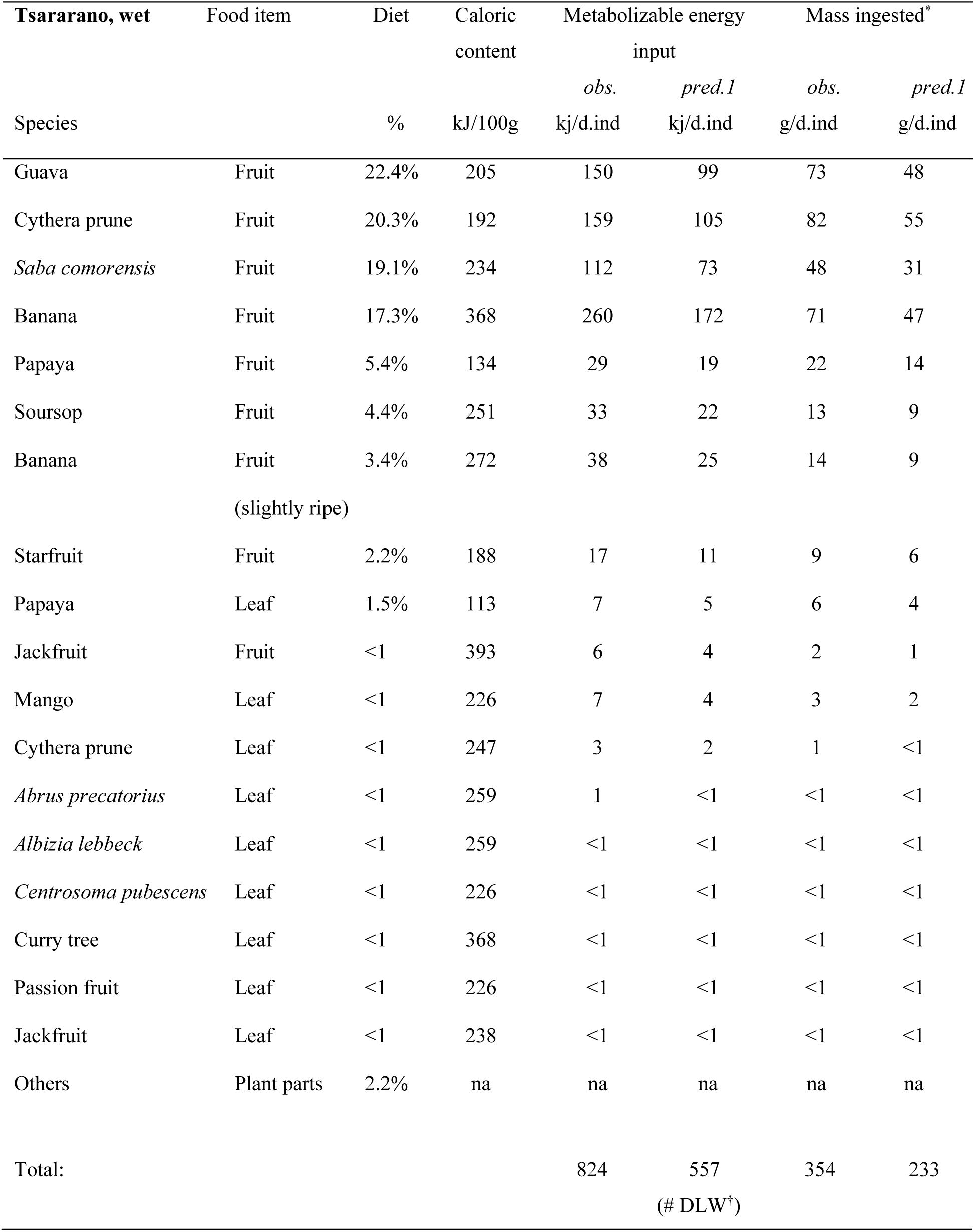
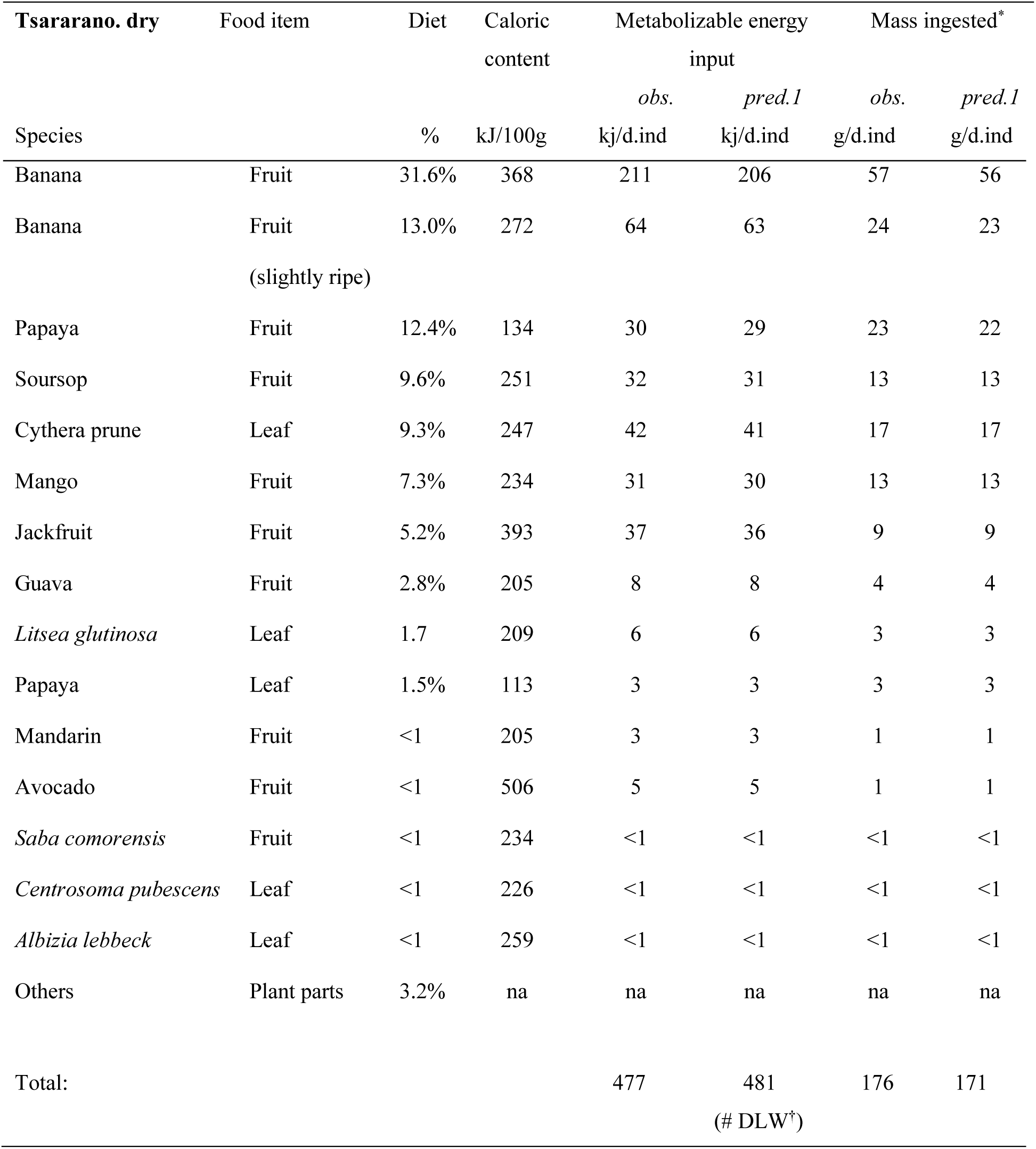
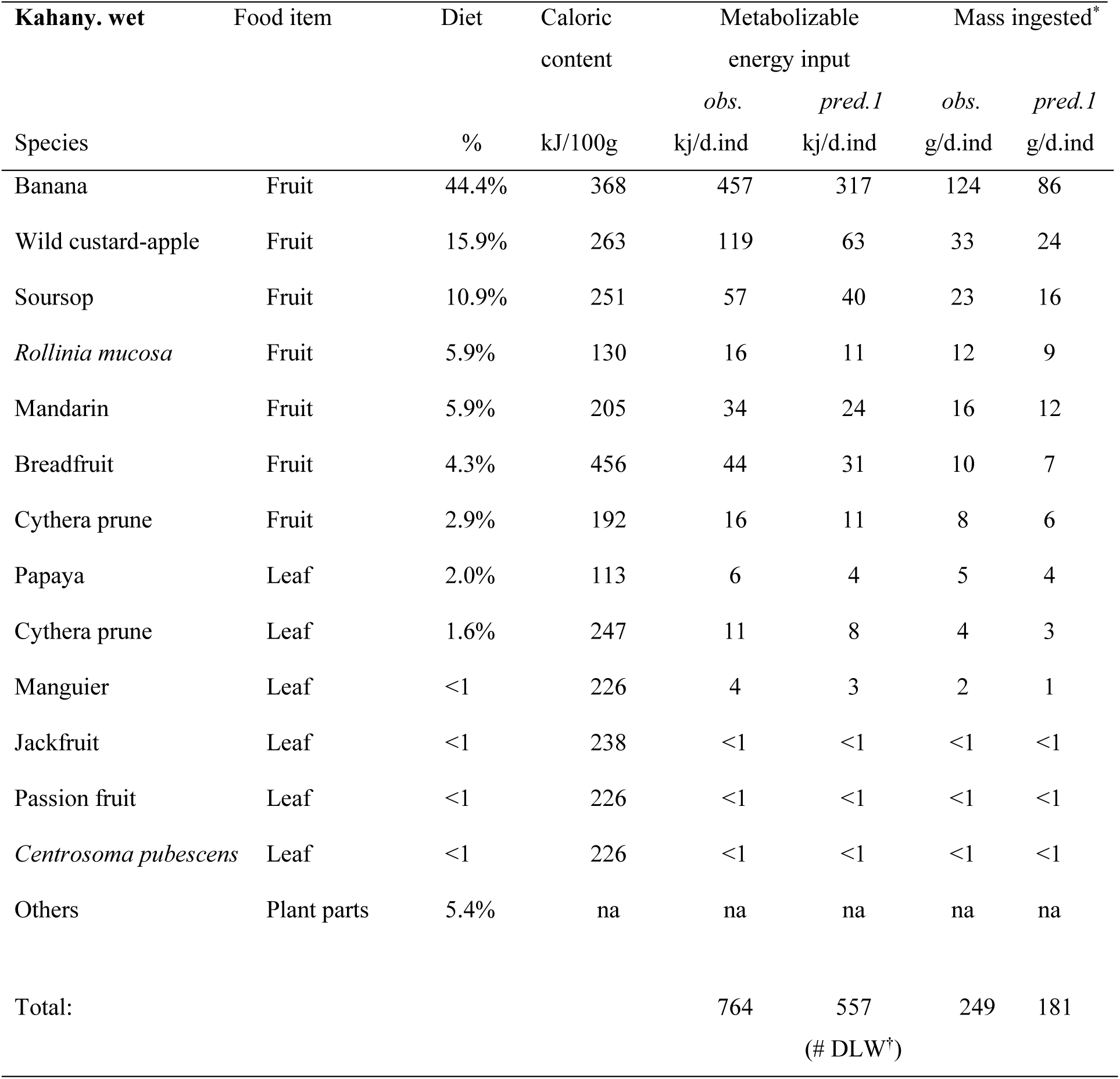
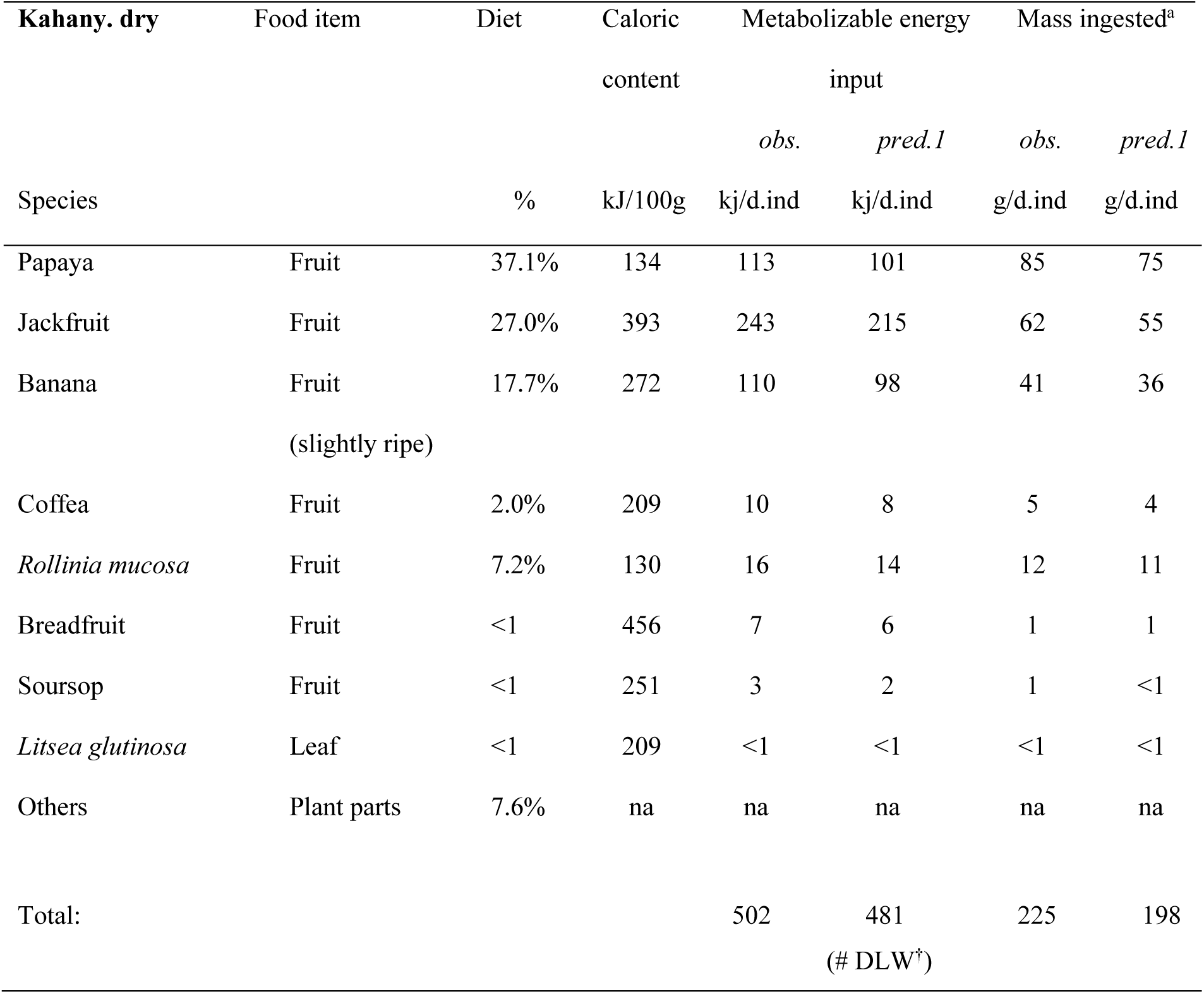
Diet and mass of food consumed by lemurs, as observed directly versus predicted by TEE (Prediction 1), at different sites and seasons in Mayotte Island. Calculations pertain to the mouthful method. obs: observed; pred: predicted; *: after swallowed seeds have been removed; ^†^: Total energy expenditure measured by doubly labelled water (DLW).

#### 3.1.3 Comparison of TEE-based predictions

The alternative predictive calculation (Prediction 2), which used the relative food proportions determined by the weighted time-sampling scores, produced similar results to Prediction 1 in terms of their correspondence with daily food intake estimated from continuous bite counts (81% ± 14% and 88% ± 21%, respectively). However, the ranking of foods in the diet differed slightly between the two, resulting in different predictions regarding the quantity of each fruit species ingested (Table S5; see Table 1 for comparison with actual food intake).

### 3.2 Testing the energetics-based method in other primates

The predicted daily food consumption (Tables S6 and S7) was, on average, 95% ± 27% (n = 10; *Eulemur rufifrons × collaris*, *Lemur catta,* and *Eulemur fulvus* in Mayotte prior to the main 2022 study; Table S9) of the amount estimated through the continuous monitoring of individual food intake (Fig. 1). Combining these results with the above predictions for *Eulemur fulvus* at Tsararano and Kahany in 2022, yielded an overall prediction accuracy of 91% ± 25% (n = 14; Table S8). In this full sample, there was no difference in the predicted and estimated food intake according to season, either in absolute value (mean difference: wet season −69 ± 39 g, dry season 12 ± 86 g; paired sample *t*-test: *t* = −1.752, *df* = 5, *p* < 0.2) or relative proportion (wet season: 80% ± 13%; dry season: 105% ± 30%; *t* = 1.635, *df* = 5, *p* < 0.2; paired sample *t*-test, Kassambara 2026; Lüdecke et al. 2021).

**Fig. 1:**
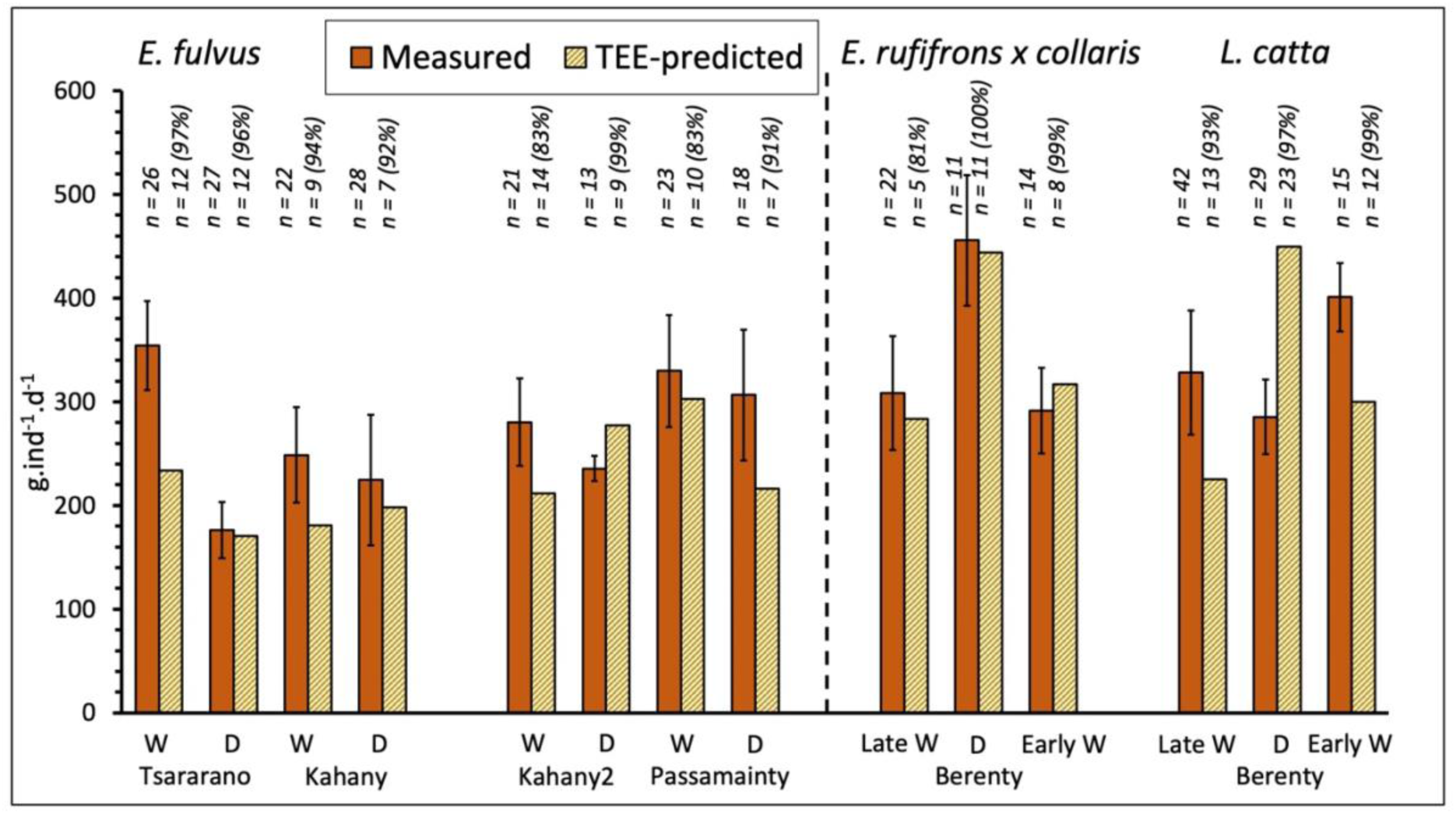
Direct estimates of, and predictions for, the daily food intake (excluding swallowed seeds) of the Mayotte lemur and other primate species, at different sites and in different seasons. The predicted figures were calculated using total energy expenditure (TEE), the proportion of food in the diet as determined by full-day continuous observations and the nutrient content of the food. The number of food items considered and their respective percentages in the diet are shown. W: wet; D: dry

### 3.3 Testing the model using limited observational data on diet

We repeated the test using dietary data limited to between one and three items, in order to simulate a situation in which the observational effort would only allow the main food items comprising a lemur’s diet to be identified – which are the ones most likely to be statistically observed. These items accounted for 43–69% of total food intake, as determined by mouthful counts. The predictive power of the TEE-based predictions was 91% ± 24% (n = 14 case studies; see Fig. S2 and Table S8).

### 3.4 Fruit loss estimates in Mayotte

Figure 2 shows the direct and predicted estimates of the mass of whole fruits (and other plant parts) collected per day by all individuals in the lemur community of each site, according to the methods used to determine or estimate food intake. Focusing on some of the major food crops selected by lemurs during the rainy season at Tsararano, the continuous bite count and TEE-based Prediction 1 revealed that an average of 7.2–10.9 kg of bananas and 7.6–11.5 kg of guavas were harvested daily (Fig. 2; Table S9). Over a two-month fruiting period from March to April, this equated to 432–656 kg of bananas and 454–690 kg of guavas. At Kahany, where lemurs are less numerous and bananas dominate the diet during the rainy season, we estimated that lemurs collected 480-660 kg of bananas during the same study period.

**Fig. 2.**
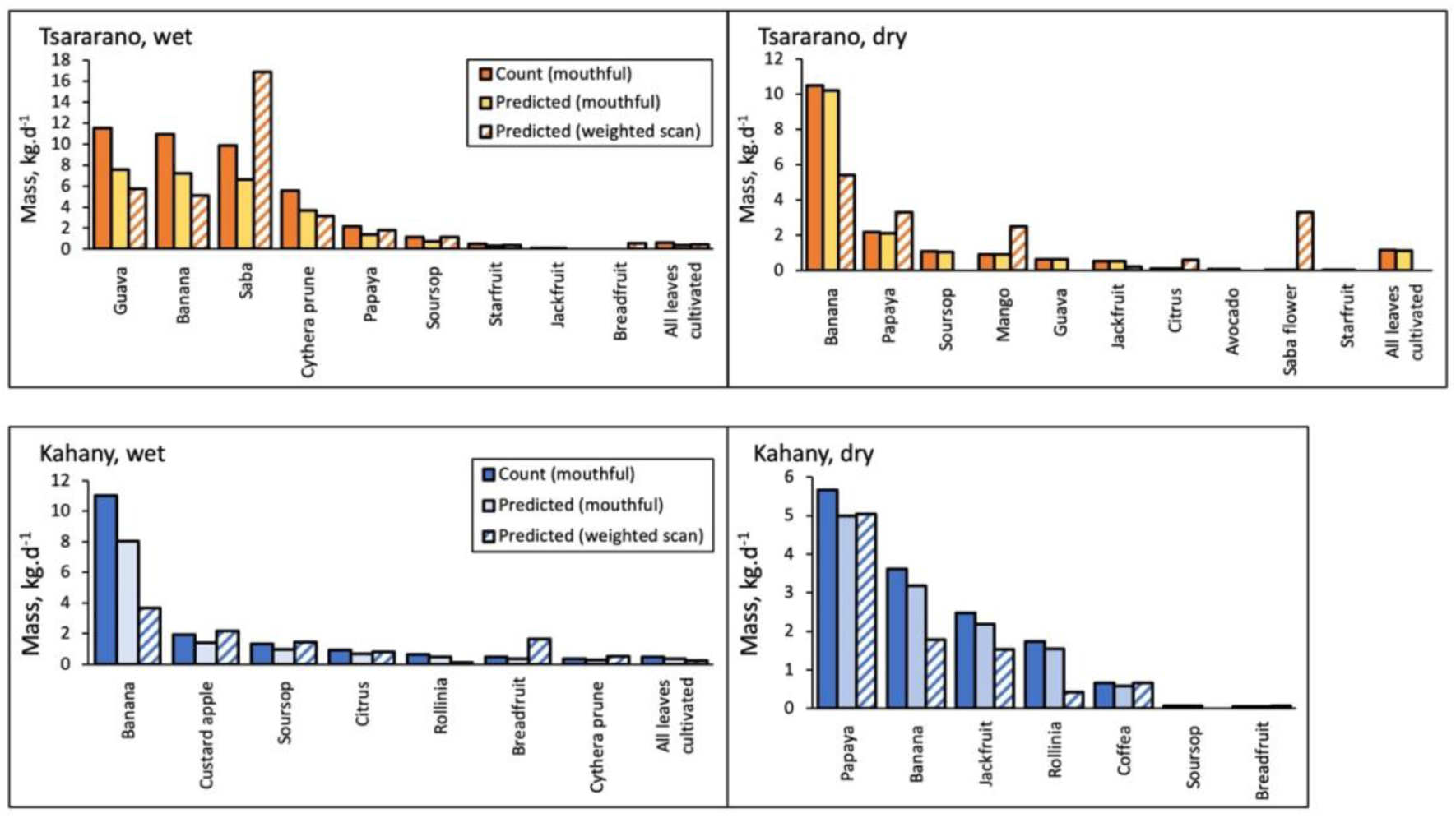
Estimated and predicted amounts of whole fruit and other plant parts removed by resident lemurs, for different plant species, at each site and in each season.

## 4 Discussion

In light of the lack of methods for assessing damage to tropical fruit crops caused by primates and other wildlife (Conforti et al., 2020), our aim was to test the feasibility of a hybrid technique primarily based on animal energy requirements for estimating crop losses. This objective was based on the premise that conservation physiology and nutritional ecology can offer integrated models for addressing scientific issues related to conservation goals in regions where humans and non-human primates coexist antagonistically (Raubenheimer et al., 2012). We demonstrate that modelling fruit production losses using an ecophysiological approach provides a reliable basis for calculating the associated economic costs.

### 4.1 TEE-based predictions

Overall, the predicted daily food intake figures (Prediction 1) were fairly close to the values estimated using the mouthful method (91% ± 25%). Apart from errors in measurement and approximations regarding the metabolizable energy content of food, some of the variability in the predictions was related to the fact that species which have evolved to cope with extreme seasonal changes do not maintain a constant energy balance. For example, although the Mayotte lemur’s daily food intake equalled the predicted amount consumed during the dry season, the lemurs ingested much more food than was required to meet their total energy needs during the rainy season (see Table 1). Similarly, *Lemur catta* in Madagascar’s natural environment consumed more food than necessary to meet their daily requirements during the rainy season, when gallery forest food resources were widely available. In both species, this reflects a period of fattening, as confirmed by body composition data (Simmen et al., 2024). *Lemur catta* consumed less food than expected during the dry season, but a detailed nutritional analysis revealed that the fat they had stored during the wet months helped them to maintain their energy balance when resources were scarce (Simmen & Rasamimanana, 2018). *Eulemur rufifrons x collaris* manages its energy budget differently to *Lemur catta*, with which it coexists, since the predictions were similar to the direct estimates of dietary intake, regardless of the season (Table S8). During the dry season, they consumed large quantities of poor-quality food, which they are able to digest despite having no identifiable anatomical digestive peculiarities (Campbel et al., 2004b; Simmen et al., 2003; Simmen & Rasamimanana, 2018). Overall, the discrepancies between the predicted and estimated values are partly connected to how the species have adapted behaviourally and physiologically to long periods of food scarcity and drought in Madagascar (Dewar & Richard, 2007; Pereira et al., 1999; Simmen et al., 2003; Wright, 1999). It is worth noting that the lemurs of Mayotte consumed different amounts of food depending on the time of year, even though they had access to a large supply of grown food crops, including during the dry season. This confirms that the periodic storage of fat in these animals is driven by internal adaptive mechanisms, influenced by photoperiod changes but contingent on food availability (Pereira et al., 1999). It is therefore important not to overlook the potential impact of a species’ energy strategy and life cycle on the quality of predictions when discussing the output of the ecophysiological model. While detailed assessments of food intake and periods of energy imbalance in primates and other mammals are scarce, further studies in this area would help to identify instances where energy intake does not match energy requirements.

To potentially simplify the application of the protocol, the results obtained using the widely adopted instantaneous-and-scan sampling method were compared with those obtained using the mouthful method. Time-derived estimates were consistent once feeding time was corrected for intake rates (Prediction 2). Staple foods remained the same, but their ranking in the diet changed however. This was to be expected because the two datasets originated from either one focal animal or a set of several group members recorded simultaneously. Furthermore, classifying food handling as part of the ‘feeding’ behavioural category while using the time sampling method artificially inflated the figures for fruit that required handling to access the edible part (e.g. *Saba*, *Arctocarpus altilis*). In this context, refining the time-sampling method to focus solely on ingestive behaviour, as with the mouthful method, would provide a more accurate representation of real ingestion.

One of the main challenges of the approach presented here is that measuring energy expenditure in the field can be difficult, invasive and expensive. Although limited, the results obtained from independent records of energy expenditure suggest that published TEE data could be used as an alternative. However, further contrast analyses in various frugivorous vertebrates may be necessary to reach a more robust conclusion. Similarly, while using a subset of dietary information to simulate incomplete knowledge of the food repertoire produced realistic estimates of daily food intake, more robust assessments of the model’s accuracy in such cases are required.

### 4.2 Fruit losses and economic impact in Mayotte

The monthly quantities of guavas and papayas harvested for sale or for processing into manufactured products at Tsararano were provided by the owner. Guavas consumed by the Tsararano lemur groups represented 42% of the production harvested between March and April (the study period). During the same period, the figure for papayas was 37%. During the dry months of the study, papayas harvested by lemurs accounted for 11% of the fruit crop. It is important to note that the owner and other farmers have reported frequent and substantial losses due to theft by humans. Large common bats also play a role, albeit a smaller one. This means that the contribution of lemurs to fruit loss, as calculated, is greatly inflated relative to their actual impact.

Based on the local consumer price index for fruits included in the basic food basket (Agreste, 2022b, c), the financial loss for the main fruits consumed by lemurs (direct food intake estimates for banana, guava, papaya, soursop, sweetsop, sugar apple, mandarin, starfruit, Cythera plum, coffee and jackfruit) is roughly equivalent to 1.17 € per lemur per day in Tsararano and 0.96 € per lemur per day in Kahany during the rainy months of the study. During the dry season, the financial loss is equivalent to 0.58 € per lemur per day for Tsararano and 0.82 € per lemur per day in Kahany. Extrapolating these figures, along with those obtained by the predictive model based on bite size, to the entire lemur population within the two sites reveals significant financial losses during the rainy months of the study (March-May; Tsararano: €811 per hectare per month; Kahany: €833 ha^-1^ month^-1^), as well as during the dry months of the study (June-July; Tsararano: €399 ha^-1^ month^-1^; Kahany: €711 ha^-1^ month^-1^). It should be noted that these crude estimates of fruit loss do not consider the lemurs’ wasteful behaviour or the damage they inadvertently cause to plant parts (e.g. papaya leaves during foraging), which can reduce the plant’s investment in fruit. They also do not include flower ingestion, which reduces fruit production. Contrasting these financial losses with the latest report on the median monthly income in Mayotte (€260 per household in 2018, with significant inequalities; INSEE, 2020) reveals that households using small fields for subsistence, or selling fruit from one-to a few-hectare fields (either raw or processed), may have experienced a significant loss of income.

The total amount of cultivated fruit lost per group varied across the different study sites in Mayotte, averaging around 1.6 kg per hectare per day (Table 2). This depended on the size of the home ranges, the time of year, and the population density of resident lemurs. These unprecedented estimates for small, complex agroecological zones, in which animals depend almost exclusively on fruit crops, could be a valuable contribution to regional-level negotiations between stakeholders. In this context, the repeated lemur censuses carried out across the entire island of Mayotte since 1974 (Tarnaud, 2024) provide supplementary data that can be used to estimate fruit losses on a larger scale.

**Table 2:**
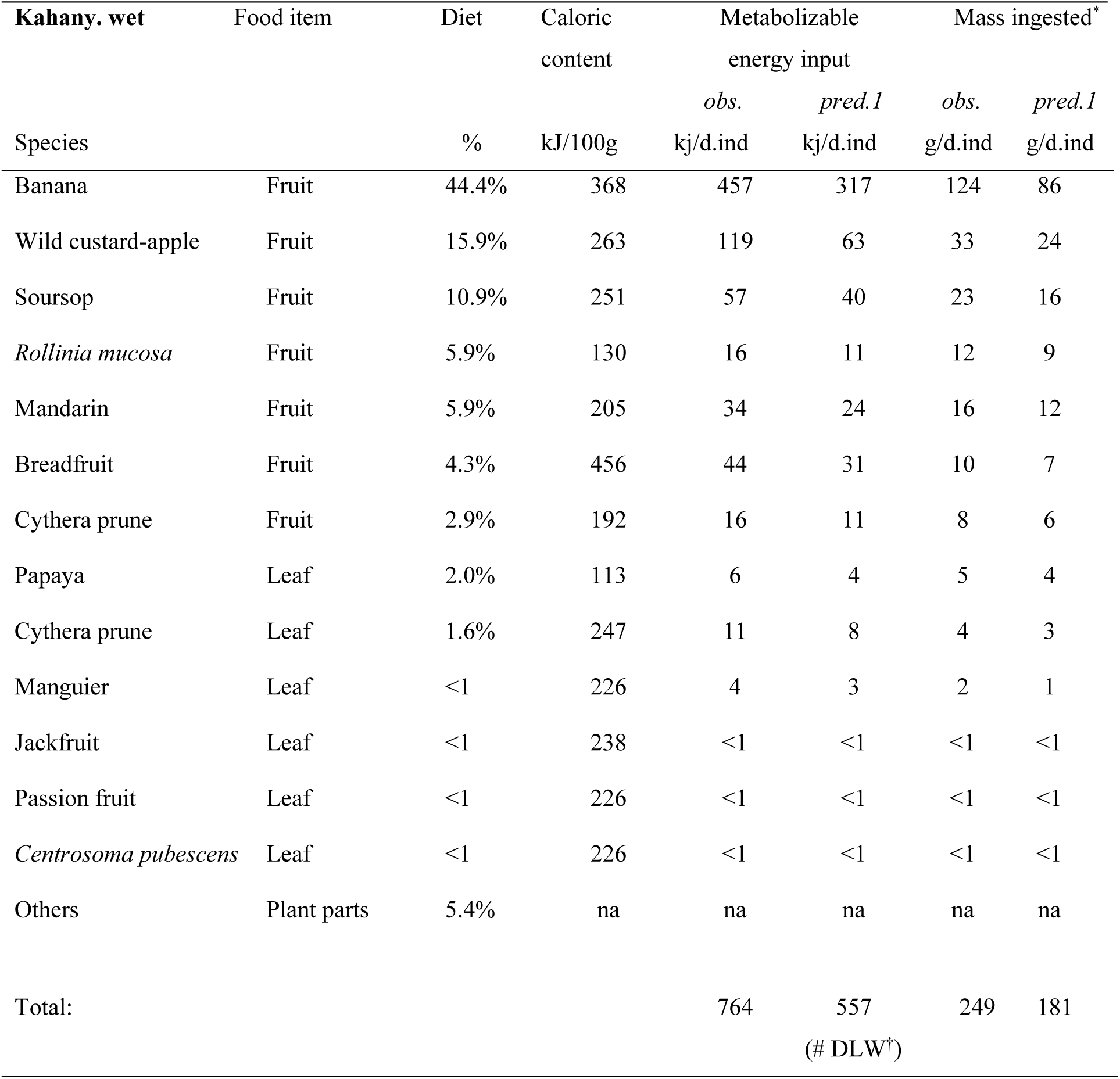
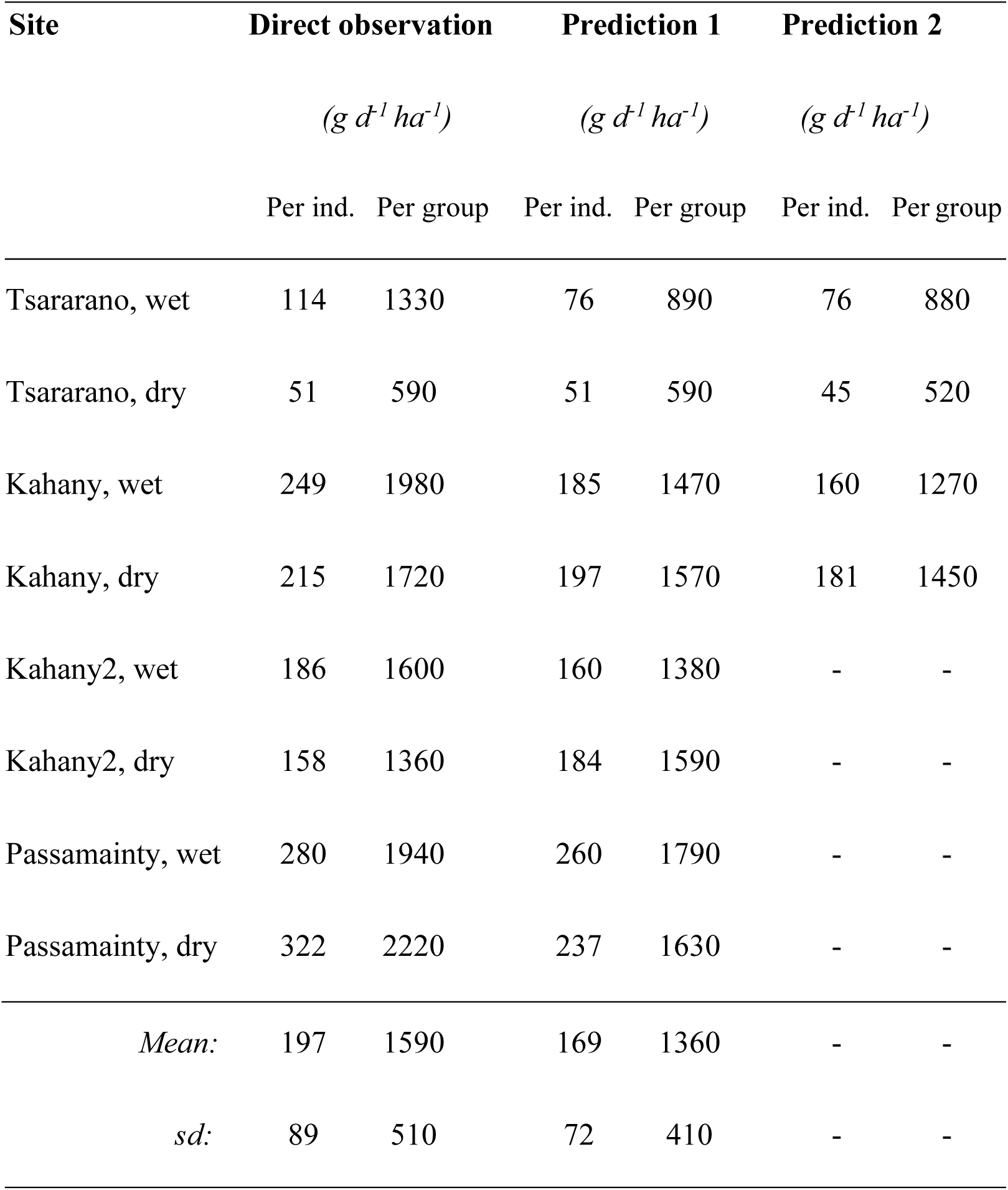
Estimates of the total daily mass (in grams) of whole cultivated fruits removed per individual (ind.) per hectare per day according to direct quantification and TEE-based predictions, for each site and season in *Eulemur fulvus*. Diet composition was assessed using either the bite size method (Prediction 1) or the weighted scan sampling method (Prediction 2).

## 5 Conclusion

The coexistence of frugivorous species alongside small-scale farmers presents a particular challenge when it comes to improving the socio-economic conditions of these small households, largely due to the inherent nature of agroforestry systems. In the context of the sustainable management of subsistence agroforests and small-scale, locally marketed crops, modelling fruit production losses caused by mammals using the proposed hybrid approach provides a quantified basis for calculating these economic losses. The accuracy of the predictions depends on the quality of the available information on the target animal, its physiology, local population density and food sources. Nevertheless, this ecophysiology method effectively provides institutional stakeholders, farming associations and NGOs with a quantitative basis for implementing financial solutions.

## Author Contributions

BS designed the study, with contributions from LT and GCP. GCP, RI, JT, BS and LT collected the data. BS wrote the first draft of the manuscript, to which all authors contributed subsequent revisions, including the final version.

## Acknowledgments

We would like to express our gratitude to the François Sommer Foundation for funding the ‘Human-Wildlife Conflicts’ project from 2021 to 2023. We would also like to thank Laurane Rami and Bernard Riera for analysing plant data in Tsararano and Kahany. This paper was enriched by discussions on payments for ecosystem services with Anouch Missirian and Sébastien Desbureaux. We are grateful to Muslim Payet and Maoulida Tchangama for granting us permission to conduct observations and experiments on their property.

## Data Availability Statement

The authors declare no conflict of interest.

