## Supplemental TableS1 for "Wildlife damage to tropical agroforests: predicting primate-induced fruit loss using species’ energy requirements"

| **Species** | **Type item** | **Moisture**  (%) | | | **Energy content**  (kJ/g) | ***Source*** | |
| --- | --- | --- | --- | --- | --- | --- | --- |
| *Abrus precatorius* | Leaf | | 79 | 259 | | | *1* |
| *Albizia spp.* | Leaf | | 80 | 259 | | | *1* |
| *Annona mucosa* | Fruit | | 92 | 130 | | | *4* |
| *Annona muricata* | Fruit | | 82.6 | 251 | | | *1* |
| *Annona squamosa* | Fruit | | 83 | 263 | | | *10* |
| *Artocarpus altilis* | Fruit | | 71.9 | 456 | | | *1* |
| *Artocarpus heterophylla* | Leaf | | 85.4 | 238 | | | *10* |
|  | Fruit | | 72.9 | 393 | | | *2* |
| *Averrhoa carambola* | Fruit | | 87.4 | 188 | | | *1* |
| *Carica papaya* | Leaf | | 75 | 113 | | | *7, 8* |
|  | Fruit | | 90.8 | 134 | | | *1* |
| *Citrus reticulata* | Fruit | | 86.1 | 205 | | | *1* |
| *Coffea sp.* | Fruit | | 74 | 209 | | | *5, 6* |
| *Hibiscus sabdariffa* | Leaf | | 85.6 | 184 | | | *1* |
| *Listea glutinosa* | Leaf | | 61.7 | 209 | | | *9* |
| *Mangifera indica* | Leaf | | 82.1 | 226 | | | *2* |
|  | Fruit | | 84.2 | 234 | | | *1* |
| *Murraya koenigi* | Leaf | | 73 | 368 | | | *2* |
| *Musa sapientum* | Fruit ripe | | 77 | 368 | | | *1* |
|  | Fruit (slightly ripe) | | 75 | 272 | | | *3* |
| *Persea americana* | Fruit | | 80.3 | 506 | | | *1* |
| *Psidium guajava* | Fruit | | 82 | 205 | | | *10* |
| *Saba comorensis* | Fruit | | 63.8 | 234 | | | *this study* |
| *Spondias dulcis* | Leaf | | 81 | 247 | | | *10* |
|  | Fruit | | 86.9 | 192 | | | *2* |

Table S1. Food energy content, expressed per unit weight wet matter
