## Supplementary figures and images for "Wildlife damage to tropical agroforests: predicting primate-induced fruit loss using species’ energy requirements"

### Supplemental FigureS1

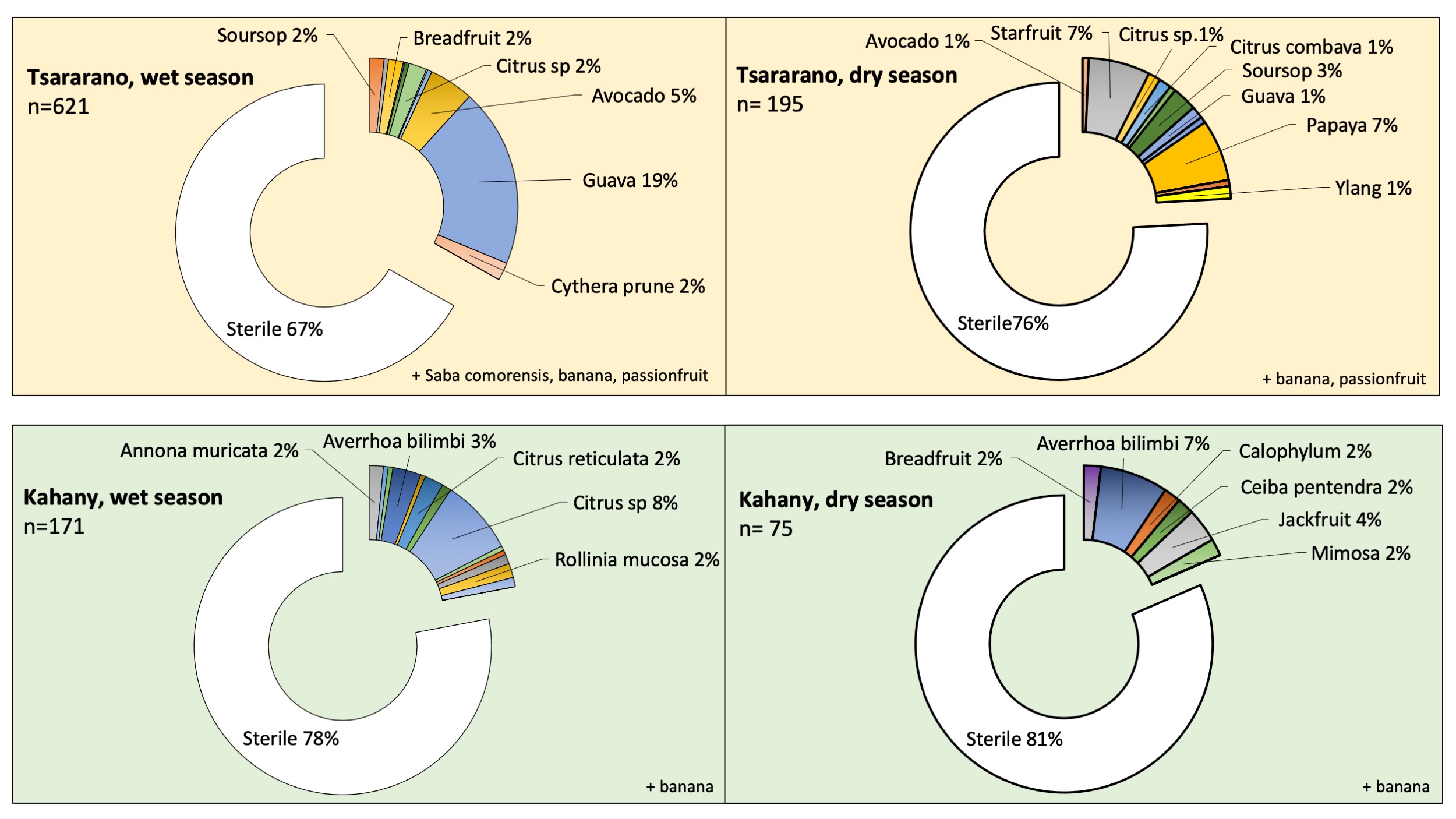

### Supplemental FigureS2

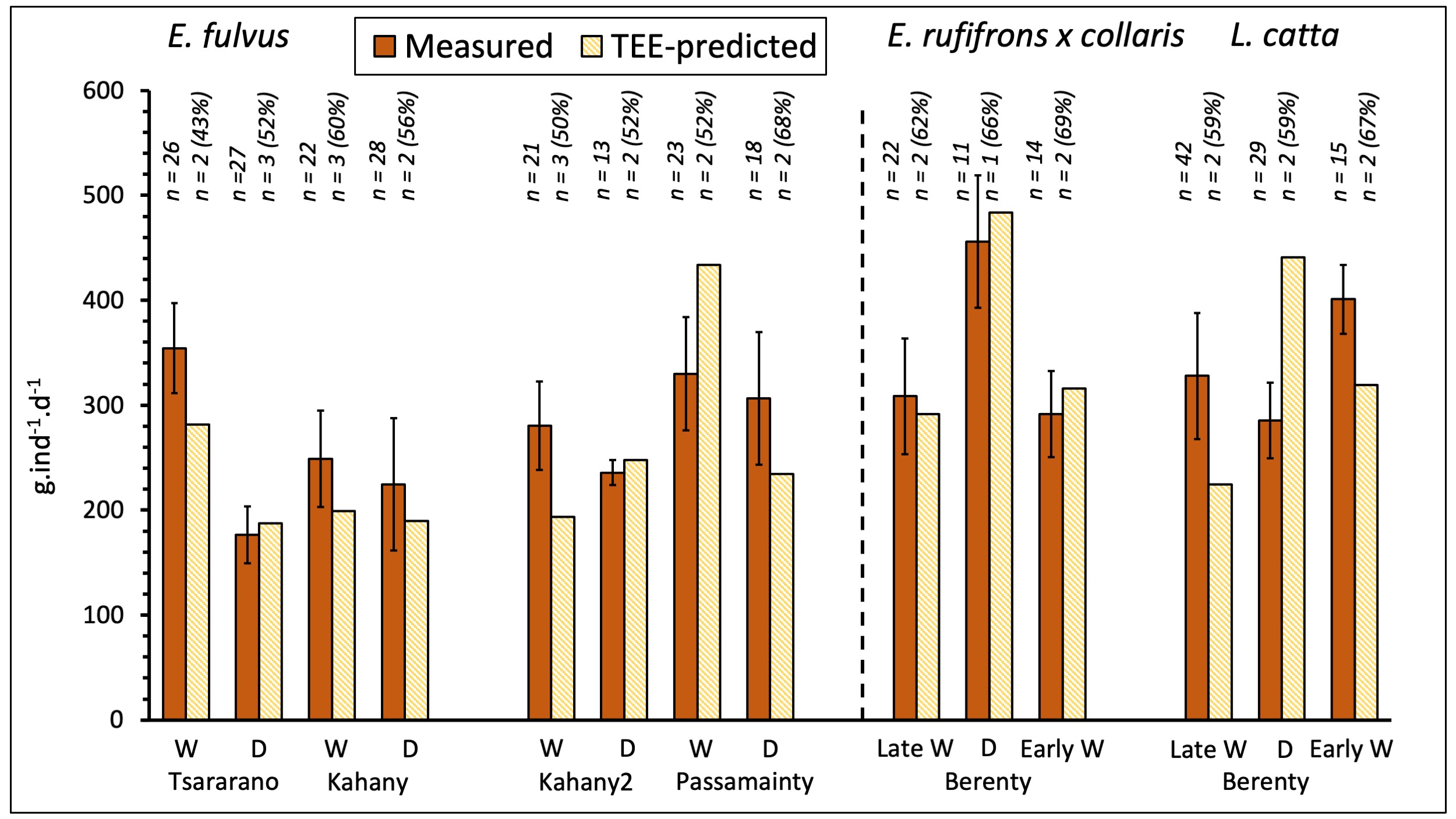
